# Abrogation of pathogenic attributes in drug resistant *Candida auris* strains by farnesol

**DOI:** 10.1101/792168

**Authors:** Vartika Srivastava, Aijaz Ahmad

## Abstract

*Candida auris*, a decade old *Candida* species, has been identified globally as a significant nosocomial multidrug resistant (MDR) pathogen responsible for causing invasive outbreaks. Biofilms and overexpression of efflux pumps such as Major Facilitator Superfamily and ATP Binding Cassette are known to cause multidrug resistance in *Candida* species, including *C. auris*. Therefore, targeting these factors may prove an effective approach to combat MDR in *C. auris*. In this study, 25 clinical isolates of *C. auris* from different hospitals of South Africa were used. All the isolates were found capable enough to form biofilms on 96-well microtiter plate that was further confirmed by MTT reduction assay. In addition, these strains have active drug efflux mechanism which was supported by rhodamine-6-G extracellular efflux and intracellular accumulation assays. Antifungal susceptibility profile of all the isolates against commonly used drugs was determined following CLSI recommended guidelines. We further studied the role of farnesol, an endogenous quorum sensing molecule, in modulating development of biofilms and drug efflux in *C. auris*. The MIC for planktonic cells ranged from 62.5-125 mM and for sessile cells was 125 mM (0 h and 4 h biofilm) and 500 mM (12 h and 24 h biofilm). Farnesol inhibited biofilm formation, blocked efflux pumps and downregulated biofilm- and efflux pump-associated genes. Modulation of *C. auris* biofilm formation and efflux pump activity by farnesol represent a promising approach for controlling life threatening infections caused by this pathogen.

## Introduction

*Candida auris* has now well evolved MDR pathogen, which has caused serious outbreaks in several continents. It was first isolated from external ear of a Japanese patient in 2009) (1) and within a decade infection caused by *C. auris* has spread rapidly across sex continents (1, 2). Centers for Disease Control and Prevention (CDC) has declared *C. auris* as a global threat with a report of causing several outbreaks in different countries, including United States (https://www.cdc.gov/fungal/candida-auris/tracking-c-auris.html). *C. auris* is causing serious bloodstream infections and other infections ranging from meningitis, bone infections, surgical wound infections and urinary tract infections have been reported in hospitals (3). *C. auris* infections are stubborn because it is resilient to the available antifungal drugs including fluconazole (first-line antifungal drug) and amphotericin B (gold standard antifungal drug). In one of the reports, CDC has analyzed antifungal susceptibility profile of different *C. auris* isolates and it was reported that almost all the isolates were resistant to azoles (fluconazole) and 1/3 of isolates remain unaffected to polyenes (amphotericin B). Whereas, echinocandins class of drugs was found active against most of the isolates of *C. auris*; however, echinocandin resistance can develop while the patient is being treated (https://www.cdc.gov/fungal/antifungal-resistance.html). Researchers have reported multidrug resistance among *C. auris* isolates as a common phenomenon, severely restraining its treatment possibilities (4). In South Africa, the first instance of infection caused by *C. auris* was reported in the 2009 and around 1,700 cases were detected between 2012 and 2016. Currently, *C. auris* is a widespread problem as it is found in almost 100 hospitals across South Africa with a vast majority of cases been reported in Gauteng (5).

Increasing prevalence of *C. auris* infection worldwide especially in South Africa motivated us to study pathogenic traits of this species. Farnesol, a first quorum sensing (QS) molecule identified in eukaryotic microorganisms (6), play an important role in an array of biological functions such as virulence, biofilm formation, and competence (7). Numerous studies have reported the effect of farnesol on *C. albicans* growth and pathogenesis (8-10). In *C. albicans*, farnesol inhibits the dimorphism (6) that prevent its establishment in different environmental conditions (11), it has antioxidant effects (12) and it inhibits transporters (13).

With this background, we emphasized to study the effect of farnesol on growth, biofilms and reversal of drug resistance in different *C. auris* isolates.

## Materials and Methods

### *Candida* isolates

In this study, 26 isolates of *Candida* spp. including 25 *C. auris* and 1 *C. albicans* SC5314 were used (**Table 1)**. All the 25 clinical *C. auris* strains were obtained from the Division of Mycology, National Institute of Communicable Diseases, Johannesburg, South Africa. The isolates were stored in glycerol stock at -80 °C until required. Farnesol, fluconazole (FLZ) and amphotericin B (AmB) were purchased from Sigma-Aldrich (USA). All the other inorganic chemicals were of analytical grade and were procured from Merck (PTY) Ltd (USA).

**Table 1:** List of *Candida* isolates.

| ***Candida* spp. isolates used in study** |  |
| --- | --- |
| *C. auris* MRL 6326 | *C. auris* MRL 3499 |
| *C. auris* MRL 6183 | *C. auris* MRL 6194 |
| *C. auris* MRL 4888 | *C. auris* MRL 6005 |
| *C. auris* MRL 6015 | *C. auris* MRL 6057 |
| *C. auris* MRL 6333 | *C. auris* MRL 5762 |
| *C. auris* MRL 4587 | *C. auris* MRL 6173 |
| *C. auris* MRL 6334 | *C. auris* MRL 5765 |
| *C. auris* MRL 3758 | *C. auris* MRL 2397 |
| *C. auris* MRL 6059 | *C. auris* MRL 5339 |
| *C. auris* MRL 4000 | *C. auris* MRL 5418 |
| *C. auris* MRL 6065 | *C. auris* MRL 6277 |
| *C. auris* MRL 2921 | *C. auris* MRL 5322 |
| *C. auris* MRL 6125 | *C. auris* MRL 6339 |
| *C. auris* MRL 6338 | *C. albicans* SC5314 |

### Antifungal susceptibility profiles

The minimum inhibitory concentration (MIC) for *C. auris* isolates against antifungal drugs was determined by broth microdilution susceptibility testing as per the recommended guidelines of Clinical and Laboratory Standards Institute (CLSI) reference document M27-A3 (14). The stock solution of AmB was prepared by using 1% dimethyl sulfoxide (DMSO) and the range of concentration tested was 16 – 0.031 µg/ml. Similarly, FLZ stock solution was prepared using deionized water and the range of concentration tested ranged from 1000 – 1.0 µg/ml. Farnesol stock solution was prepared with 1% DMSO and the concentration tested ranged from 250 – 0.48 mM. In every set of experiment, cell free (sterility) and drug free (growth) controls were included for each *C. auris* isolates and all the isolates were tested in triplicate. *C. albicans* SC5314 was kept as a standard laboratory control in each test performed. The MICs were defined as the lowest concentration of drugs or farnesol that resulted in the complete inhibition of growth.

### *Candida auris* biofilm formation

Biofilm formation by *Candida* spp. on medical devices is very common problem and life threatening for patients. Different clades of *C. auris* have also been reported to produce biofilm and therefore we studied the biofilm forming capability of South African clade *C. auris* isolates. The method adapted to evaluate biofilm formation by *C. auris* isolates has been described previously (15). The metabolic activity of *C. auris* biofilm was also compared with *C. albicans* SC5314 biofilms.

### Effect of farnesol on sessile *C. auris* cells and successive development of biofilms

To evaluate the activity of farnesol on sessile cells of *C. auris* and latter development of biofilms, a method described previously was followed (15). Briefly, 100 µl standardized cell suspensions (5.0 × 10^6^ CFU/ml) were inoculated into predetermined wells of 96 well microtiter plates followed by incubation at 37 °C for 0, and 4 h. After incubation the growth medium was removed followed by thorough washing with sterile Phosphate Buffer Saline (PBS). After removal of non-adherent cells, different concentrations (500 mM to 0.488 mM) of farnesol were added to the wells of microtiter plate. To check the effect of farnesol on adherence and biofilm formation, farnesol and standardized suspension were added together to microtiter plate (zero time/preincubation) and incubated for 48 h. Furthermore, to see the effect on 4 h mature biofilms, cells were incubated under biofilm forming conditions for 4 h and then sessile cells were removed, washed gently with sterile PBS and farnesol was added to predetermine wells and incubated for 48 h. Metabolic activities of the biofilms were measured using a 3-(4,5-Dimethyl-2-thiazolyl)-2,5-diphenyl-2H-tetrazolium bromide (MTT) reduction assay, as previously described (15).

### Effect of farnesol on mature biofilms

*C. auris* biofilms were allowed to grow for 12 and 24 h at 37 °C under favorable biofilm forming conditions. The growth medium was removed and biofilm was washed gently with sterile PBS. Farnesol (500 mM to 0.488 mM) was added to the predefined wells of microtiter plates and further incubated for 24 h at 37 °C. The metabolic activity of treated and untreated biofilms was assessed by MTT reduction assay. Biofilm inhibitory concentrations (BIC) were defined as the lowest concentration of farnesol where we report inhibition (≥ 90%) compared to the growth control.

### Confocal laser scanning microscopy (CLSM)

To further confirm the effect of farnesol on *C. auris* biofilm, CLSM was done. *C. auris* strain MRL 5765 was allowed to grow on glass coverslips in 6-well microtiter plates under biofilm forming conditions. Farnesol (BIC) was administered in designated wells at different time points (4 h, 12 h and 24 h) except the growth control wells (untreated cells). The plates were further incubated for 24 h at 37 °C. Following incubation, the planktonic cells were aspirated and biofilms were gently washed twice with PBS and stained with fluorescent dye FUN-1 (Invitrogen, Thermo Fisher Scientific, ZA) and concanavalin A (ConA)-Alexa Fluor 488 conjugate (Invitrogen, Thermo Fisher Scientific, ZA). For staining, the coverslips were transferred to a new 6-well microtiter plate and incubated with 2 ml PBS containing FUN-1 (10 µM) and ConA-Alexa Fluor 488 conjugate (25 µg/ml) for 45 min at 37 °C in dark. FUN-1 (excitation wavelength = 543 nm and emission wavelength = 560 nm) is a vital dye and only live cells are capable of transporting it to the vacuole and result into orange-red cylindrical intra-vacuolar structures (CIVS) whereas in dead cells FUN-1 remain in the cytosol and fluoresces yellow-green (16). ConA (excitation wavelength = 488 nm and emission wavelength = 505 nm) on the other hand fluoresces bright green when binds to α-mannopyranosyl and α-glucopyranosyl residues present in cell wall and biofilm matrix. After incubation with fluorescent dyes the glass coverslips were flipped on glass plates and stained biofilms were observed using a Zeiss Laser Scanning Confocal Microscope (LSM) 780 and Airyscan (Carl Zeiss, Inc.). Multitrack mode was used to collect the images of green (ConA) and red (FUN-1) fluorescence simultaneously. The thickness or volume of whole biofilm was determined by collecting Z-stack picture and the distances between first and last fluorescent confocal plane was defined as biofilm thickness (17).

### Extracellular Rhodamine 6G efflux assay

Extracellular efflux of Rhodamine 6G (R6G) from *C. auris* cells were evaluated as described previously (18) with some minor adjustments. For this study four *C. auris* isolates (MRL 2397, MRL 5765, MRL 5762, and MRL 6057) were selected and *C. albicans* SC5314 was used as standard for efflux activity. Briefly, *Candida* cells were grown on Sabouraud Dextrose Agar (SDA) plates for 24 h at 37 °C. The cells (5.0 × 10^6^ CFU/ml) were inoculated in 50 ml growth media (SDB) for 8 h at 37 °C. Post incubation media was centrifuged (3000 rpm), washed with 25 ml PBS (without glucose) at least two times. The washed cells were resuspended in sterile glucose-free PBS (2% cell suspension). The cells were further incubated in 50 ml PBS containing 2-deoxy-D-glucose (5.0 mM) and 2,4 dinitrophenol (5.0 mM) for 45 min, resulting in de-energizing of cells. Followed by de-energization the cells were washed and again resuspended in glucose-free PBS (2% cell suspension), R6G (final concentration of 10 μM) was added to this resuspension and incubated for 40 min at 37 °C. After incubation cells were again washed and resuspended in glucose-free PBS (2% cell suspension). Samples (2 ml) were withdrawn at definite intervals (0, 5, 10, 15, 20 min). After harvesting samples were pelleted at 3,000 rpm and optical density of supernatant was recorded at 527 nm. To study the energy dependent R6G efflux, glucose (0.1 M) was added after 20 min incubation to the cells resuspended in glucose-free PBS. The absorbance was recorded till 60 min of incubation with glucose and the last reading was recorded after overnight (20 h; 1200 min) incubation. Positive as well as negative controls were included in all the experiments. The standard concentration curve of R6G was prepared for determining the actual concentration of R6G effluxed.

For competition assays, yeast cells were exposed for 2 h to different concentration of farnesol (0.5 × MIC and MIC). Post exposure the cells were pelleted (3000 rpm) and washed twice with sterile PBS (without glucose). Thereafter treated cells were de-energized and then equilibrated in R6G as stated above. Samples (2 ml) were withdrawn at predetermined time points (0, 5, 10, 15, 20 min), centrifuged (3,000 rpm) and absorbance of supernatant was recorded at 527 nm. The estimation of energy dependent R6G efflux was done by adding glucose (0.1 M) after 20 min incubation to the resuspended cells and reading were recorded till 60 min and last reading was recorded after 1200 min of incubation. Positive as well as negative controls were included in all the experiments. The standard concentration curve of R6G was prepared for determining the actual concentration of R6G effluxed.

### Intracellular Rhodamine 6G accumulation assay

Intracellular accumulation assay was executed as discussed earlier with minor modifications (18). Briefly, *C. auris* isolates (MRL 6057, MRL 2397, MRL 5762, and MRL 5765) cells were grown overnight in SDB medium at 37 °C. After incubation cells were centrifuged and washed twice in sterile PBS and re-inoculated in sterile SDB broth supplemented with farnesol (at 0.5 × MIC and MIC) for 2 h at 37 °C. Post incubation, cells were pelleted (3000 rpm) and given sterile PBS wash. The washed cells (5.0 × 10^6^ CFU/ml) were resuspended in sterile PBS (1.0 ml) supplemented with glucose (2%) and R6G (4 μM) and then incubated for 30 min at 37 °C. Post incubation cells were washed twice with cold sterile PBS and the pellet was used for fluorescence microscopy.

### Real Time PCR

The mRNA expression level for CDR1, CDR2, SNQ2, HYR3, IFF4, PGA7, PGA26, PGA52, MDR1, MDR2 and ACT1 (housekeeping gene) were measured using RT-qPCR. The method was taken from previous study described elsewhere with minor adjustments (19). Briefly, *C. auris* isolates (MRL 6057, MRL 2397, MRL 5762 and MRL 5765) were incubated overnight at 37 °C SDB medium. Overnight cultures were centrifuged (3000 rpm) resulting a pellet, which was resuspended in sterile SDB broth (10 ml) containing farnesol (MIC) and then incubated at 37 °C for 2 h. After exposure time centrifugation (3000 rpm) was done, supernatant was discarded and pellet was used for RNA extraction. RNA was extracted by using RNA MiniPrep kit (Inqaba Biotechnical Industries (Pty) Ltd) as instructed by the manufacturer. The Nano Drop 2000 plate reader (Thermo Scientific) was used to determine concentrations of isolated RNAs. Purity of RNAs was assessed by determining A_260_/A_280_ ratio and a ratio above 2 was used for further experiment (qRT-PCR). cDNA was synthesized using Lasec SA (Pty) Ltd cDNA kit following manufacturer’s instructions. PCR master mix and PowerUp SYBR Green Mater Mix (Applied Biosystems) were used to amplify *C. auris* genes from cDNA by Light Cycler Nano Real-Time PCR system (Roche). **Table 2** enlists the primers (both forward and reverse) used for amplification and experimental conditions were as follows: UDG activation at 50 °C for 2 min (Hold), Dual-lock DNA polymerase at 95 °C for 2 min (Hold), 40 cycles of denaturation at 95°C for 15 sec, annealing at 53 °C for 15 sec, and extension 72 °C for 1 min. Dissociation curve conditions (melt curve stage) were as follows: Pre-melting at ramp rate of 1.6 °C/sec, 95 °C and 15 sec; Melting at ramp rate of 1.6 °C/sec, 60 °C and 1 min; Melting at ramp rate of 0.15 °C/sec, 95 °C and 15 sec. The dissociation curve and CT values were determined using the Light Cycler Nano system. The gene expression was quantified and analyzed with respect to the housekeeping gene ACT1 using formula 2^-ΔΔCT^. The relative change in expression was estimated by normalizing to housekeeping gene (ACT1).

**Table 2:**
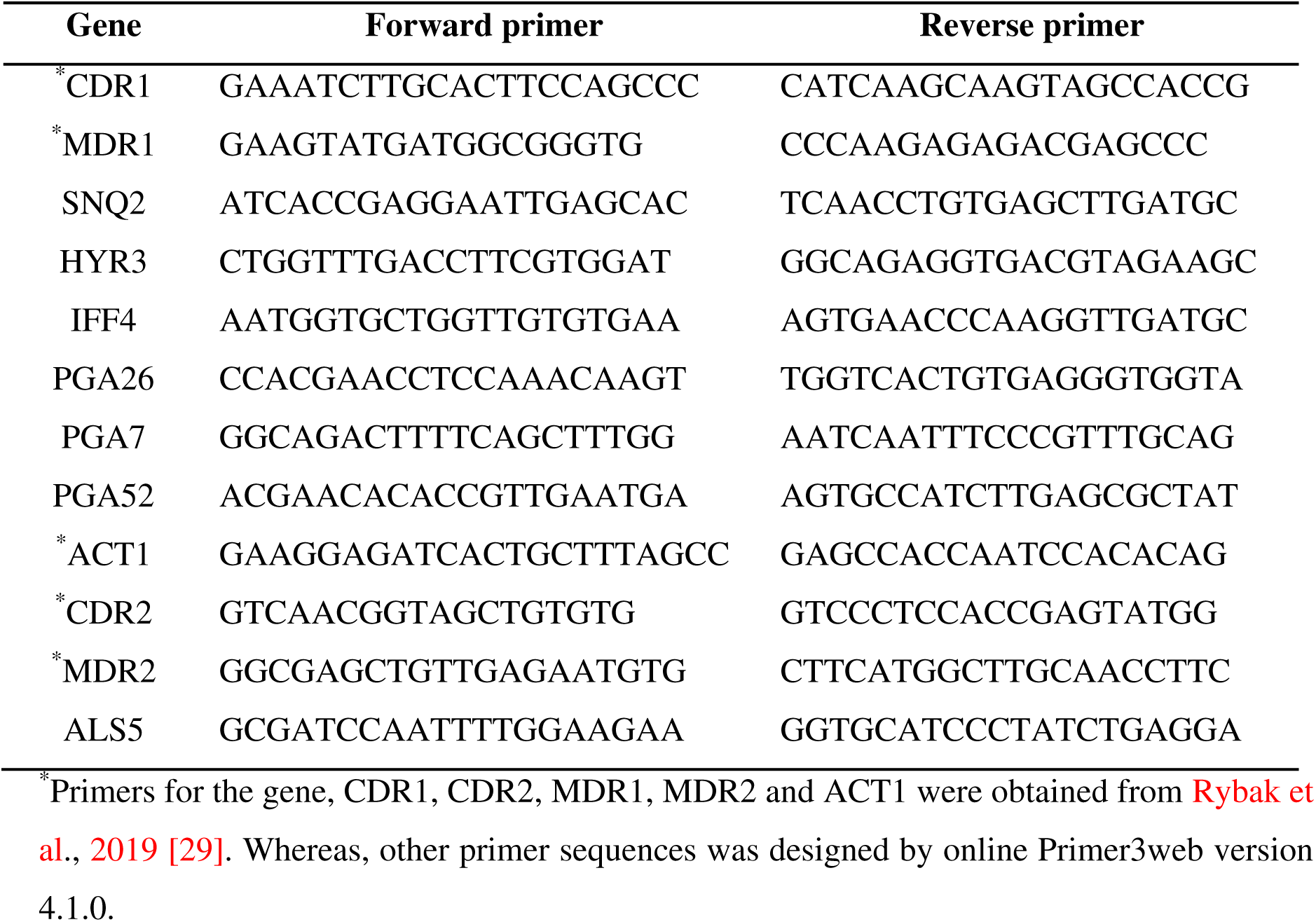
Nucleotide sequences for primers (5′—3′)

### Statistics

All the data and graphs were made and statistically analyzed using GraphPad prism version 5.01. All the experiments were carried out in triplicates, and the data obtained were presented as means ± standard error of the mean. Two-way ANOVA was used to compare untreated control with treated groups and P value less than 0.05 was considered significant.

## Results and Discussion

### Antifungal susceptibility testing

All the clinical isolates of *C. auris* used in the present study were found sensitive to the farnesol within the MIC range of 62.5 - 125 mM. MIC values for AmB ranged from 0.125 – 4.0 µg/ml whereas for FLZ the MIC values ranged from 16 – 500 µg/ml (**Table 3**). As there are no confined cutoff values to differentiate susceptible and resistant *C. auris* isolates against these drugs, it would have been inappropriate to categorize these isolates. However, CDC has established arbitrary breakpoints for *C. auris*, which were set at ≥ 32 μg/ml and ≥ 2 μg/ml for FLZ and AmB, respectively (4, 20). Based on these cutoff values, all the tested *C. auris* isolates except three (MRL 3758, MRL 2397, MRL 5322) were FLZ resistant whereas five *C. auris* isolates (MRL 2921, MRL 4000, MRL 5765, MRL 5762 and MRL 6057) were found resistant to AmB. Recent studies have also confirmed that *C. auris* isolates are usually resistant or less susceptible to azoles (21-23). Furthermore, lower susceptibility of *C. auris* isolates against AmB is also in agreement with previous studies, where high AmB MICs for *C. auris* isolates was reported (24-26). Inhibitory and modulatory effects of farnesol in *C. albicans* and other non-albicans has already being studied and its impact on biofilm formation, efflux pumps, and other virulence attributes is well established (8, 27-29). However, this study for the first time reported inhibitory effect of farnesol on *C. auris* isolates.

**Table 3:**
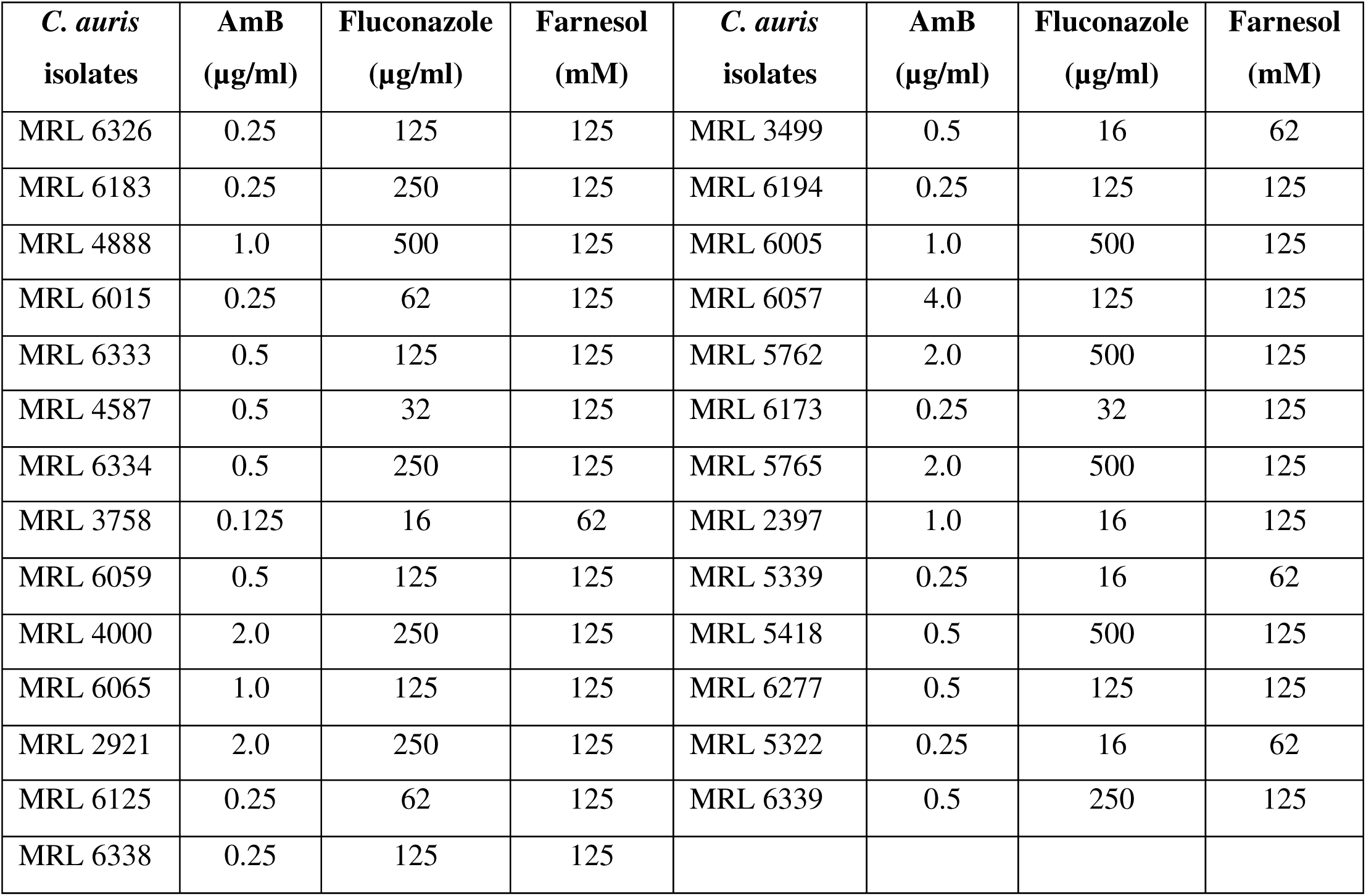
MIC values for AmB, FLZ and farnesol against isolates of *C. auris*.

| *C. auris*<br>isolates | AmB<br>(µg/ml) | Fluconazole<br>(µg/ml) | Farnesol<br>(mM) | *C. auris*<br>isolates | AmB<br>(µg/ml) | Fluconazole<br>(µg/ml) | Farnesol<br>(mM) |
| --- | --- | --- | --- | --- | --- | --- | --- |
| MRL 6326 | 0.25 | 125 | 125 | MRL 3499 | 0.5 | 16 | 62 |
| MRL 6183 | 0.25 | 250 | 125 | MRL 6194 | 0.25 | 125 | 125 |
| MRL 4888 | 1.0 | 500 | 125 | MRL 6005 | 1.0 | 500 | 125 |
| MRL 6015 | 0.25 | 62 | 125 | MRL 6057 | 4.0 | 125 | 125 |
| MRL 6333 | 0.5 | 125 | 125 | MRL 5762 | 2.0 | 500 | 125 |
| MRL 4587 | 0.5 | 32 | 125 | MRL 6173 | 0.25 | 32 | 125 |
| MRL 6334 | 0.5 | 250 | 125 | MRL 5765 | 2.0 | 500 | 125 |
| MRL 3758 | 0.125 | 16 | 62 | MRL 2397 | 1.0 | 16 | 125 |
| MRL 6059 | 0.5 | 125 | 125 | MRL 5339 | 0.25 | 16 | 62 |
| MRL 4000 | 2.0 | 250 | 125 | MRL 5418 | 0.5 | 500 | 125 |
| MRL 6065 | 1.0 | 125 | 125 | MRL 6277 | 0.5 | 125 | 125 |
| MRL 2921 | 2.0 | 250 | 125 | MRL 5322 | 0.25 | 16 | 62 |
| MRL 6125 | 0.25 | 62 | 125 | MRL 6339 | 0.5 | 250 | 125 |
| MRL 6338 | 0.25 | 125 | 125 |  |  |  |  |

### *C. auris* biofilm formation

Metabolic activity of untreated healthy biofilms was quantified by MTT and the results revealed that all the 25 isolates of *C. auris* were able to form biofilm. Whereas the extent of biofilm formed varied among the isolates, only 16 out of 25 *C. auris* isolates were good biofilm formers which was evident from higher MTT readings. Furthermore, biofilm formed by these 16 isolates were compared with biofilm formed by *C. albicans* SC5314. The metabolic activity recorded for *C. albicans* biofilm was 1.5 to 2.0 times higher than metabolic activity recorded for *C. auris* biofilm. Therefore, we concluded that *C. auris* isolates were having biofilm forming capability but less than *C. albicans* SC5314. Our results are in agreement with previous findings, which characterized and compared the biofilm formation of 16 different *C. auris* isolates with biofilms formed by *C. albicans* (23, 30). In these studies, they reported *C. auris* biofilms are mainly composed of yeast cells whereas, extremely heterogeneous architecture was found in *C. albicans* biofilms. They also investigated and compared the virulence factors of *C. auris* isolates with *C. albicans*. Researchers also described and differentiated the biofilm forming ability of non-aggregative and aggregative strains of *C. auris* (30).

### Farnesol inhibits *C. auris* biofilm development and formation

Sixteen *C. auris* isolates showing good biofilm forming capability were selected for this study. Adherent cell population (0, 4, 12 and 24 h) of *C. auris* was treated with different concentrations of farnesol (500 to 0.48 mM) to determine its inhibitory effect over *C. auris* biofilm formation. The results demonstrated that pre-incubation (0 h) of isolates with the farnesol (125 mM) stopped the adherent cells from growing further, resulting in scarce or missing biofilms in the microtiter plate. Additionally, at concentration of 7.81 mM of farnesol we observed more than 50% inhibition in biofilm formation. When the cells were allowed to grow for 4 h under biofilm forming conditions the biofilm inhibitory concentration (BIC) was found to be 125 mM. Furthermore, 12 and 24 h mature biofilm became highly resistant to low concentration of farnesol and were inhibited at a high concentration of 500 mM. This was reflected clearly in the readings recorded at 490 nm, with lowest MTT readings at the highest concentration of farnesol. Percent inhibition of *C. auris* biofilms at different concentration is shown in Fig. 1. The effect of farnesol on subsequent biofilm development decreased, as we allowed cells to adhere by increasing the incubation time. In comparison, good biofilm architecture was recorded for untreated biofilms (negative control), which was marked by readings obtained at 490 nm. The results further suggested that mature biofilms (12 and 24 h) are more resistant. Although *C. auris* forms significantly reduced biofilms as compared to *C. albicans*, nevertheless, it has the ability to form biofilms on various medical devices (31). Antifungal resistance and lower susceptibility among *C. auris* biofilms against commonly used drugs in comparison to *C. albicans* and *C. glabrata* has already been reported by researchers (30, 32). Results in the present study established the fact that farnesol prevented adherence of *C. auris* cells, which is a crucial step in biofilm formation.

**Fig. 1:**
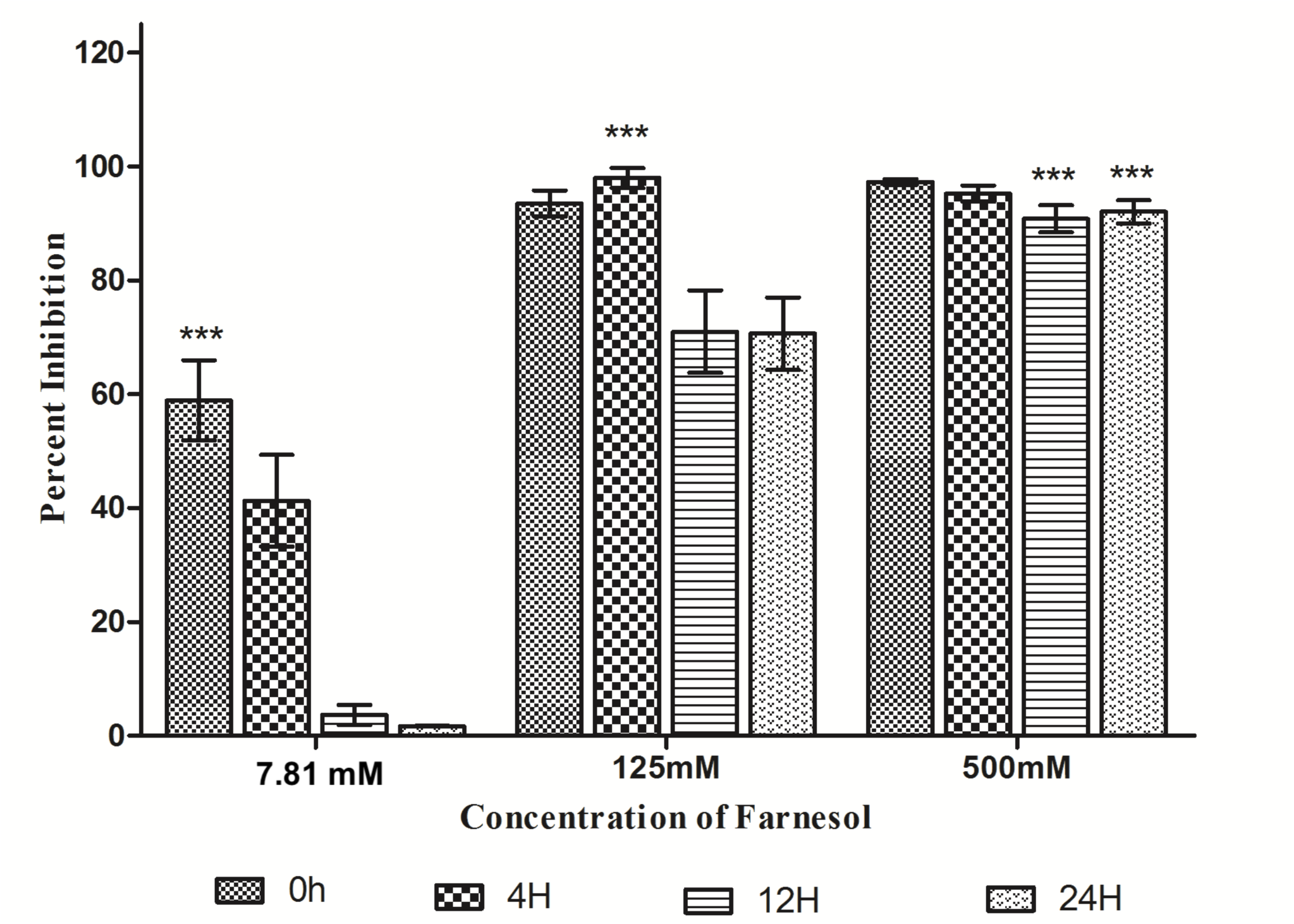
Effect of farnesol on formation of *C. auris* biofilms. *C. auris* cells were incubated under biofilm forming conditions with different concentrations of farnesol (500-0.48 mM) at time interval (0 h, 4 h, 12 h, and 24 h). Pre-incubation with farnesol (0 h) and treatment of the *C. auris* isolates with farnesol after 4 h, 12 h and 24 h incubation. MTT reduction assay was used for quantification. Two-way ANOVA was used to study statistical significance (***P < 0.001).

### CLSM analysis

*C. auris* biofilm architecture and antifungal activity of farnesol was examined by using CLSM technique. Biofilm cells at different growth phases were challenged with farnesol for 24 h and stained with FUN-1 and ConA to evaluate the cell viability and thickness of biofilm extracellular matrix. The inhibitory effect of farnesol was assessed by comparing treated biofilms with untreated control under confocal microscope. CLSM analysis of untreated control revealed that as biofilm gets more mature their metabolic activity decreases, as seen by Fun-1 staining. Fun-1 permeabilizes into cells and only metabolically active cells convert Fun-1 to orange-red CIVS (16). Highest metabolic activity was observed in 4 h biofilms that was estimated by staining intensity of CIVS (orange-red colour) whereas as the staining decreased with increase in incubation time of biofilms [Fig. 2]. After 24 h of incubation under biofilm forming conditions, only a small subpopulation showed formation of CIVS, which was evident from orange-red fluorescence. These results are incongruent with the previous findings, where cells embedded in mature biofilm (48 h) were reported to have reduced metabolic activity and limited growth as compared to cells with young biofilms (4 h) (33). Contrary to this with increasing age and maturity the thickness of extracellular matrix was found to increase which was evident by more green fluorescence during CLSM as a result of binding of ConA to glucose and mannose residues present in cell wall and biofilm matrix. The thickness of 4 h young, 12 h and 24 h mature biofilm was found to be around 9.08 µm, 12.11 µm and 13.93 µm respectively.

**Fig: 2.**
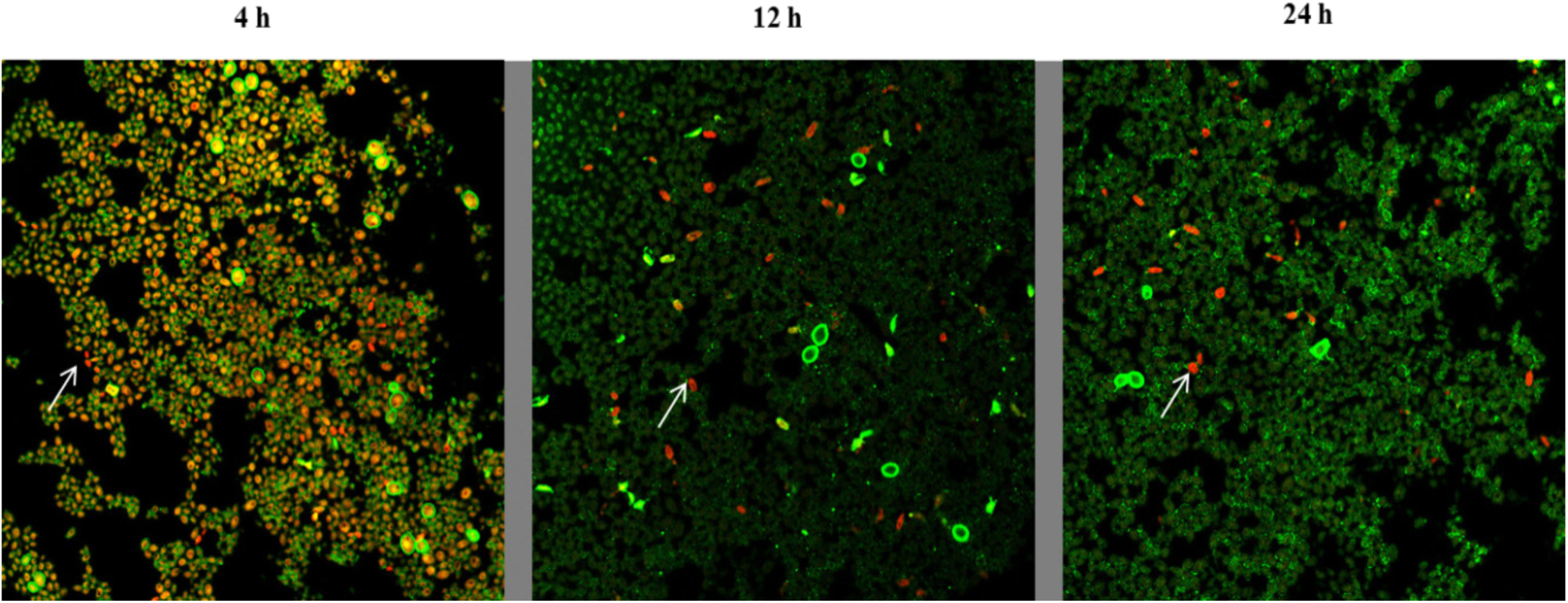
Metabolic activity of *C. auris* cells in biofilm decreases with increasing maturity. *C. auris* strain MRL 5765 cells were stained with FUN-1 and ConA-Alexa Fluor 488 conjugate stain after incubation of biofilm for 4 h, 12 h and 24 h respectively. The image utilizes a 63X oil immersion objective with magnification, X1. The cells that were live and active produce orange-red cylindrical intra-vacuolar structures (CIVS). The increase in green fluorescence shows the increase in biofilm extracellular matrix with increasing incubation time.

In our current study, CLSM revealed that after treatment with farnesol (BIC; 125 mM for 4 h and 500 mM for 12 h and 24 h) the biofilms were abrogated and the viability of cells in the biofilm was altered, which was evident by aggregate of cells scattered on the coverslip and presence of yellow-green structures showing metabolically inactive cells (Fig. 3). Furthermore, farnesol also decrease the whole volume of the biofilm, the thickness of treated 4 h, 12 h, and 24 h biofilms were measured and found to be 8.48 µm in case of treated 4 h, and 12 h biofilms whereas the thickness of treated 24 h biofilm was recorded as 9.69 µm. It is clear that farnesol significantly affected the sustainability of yeast cells in biofilms pointing towards the ability of farnesol to penetrate cell membrane resulting in an effective anti-biofilm activity.

**Fig: 3.**
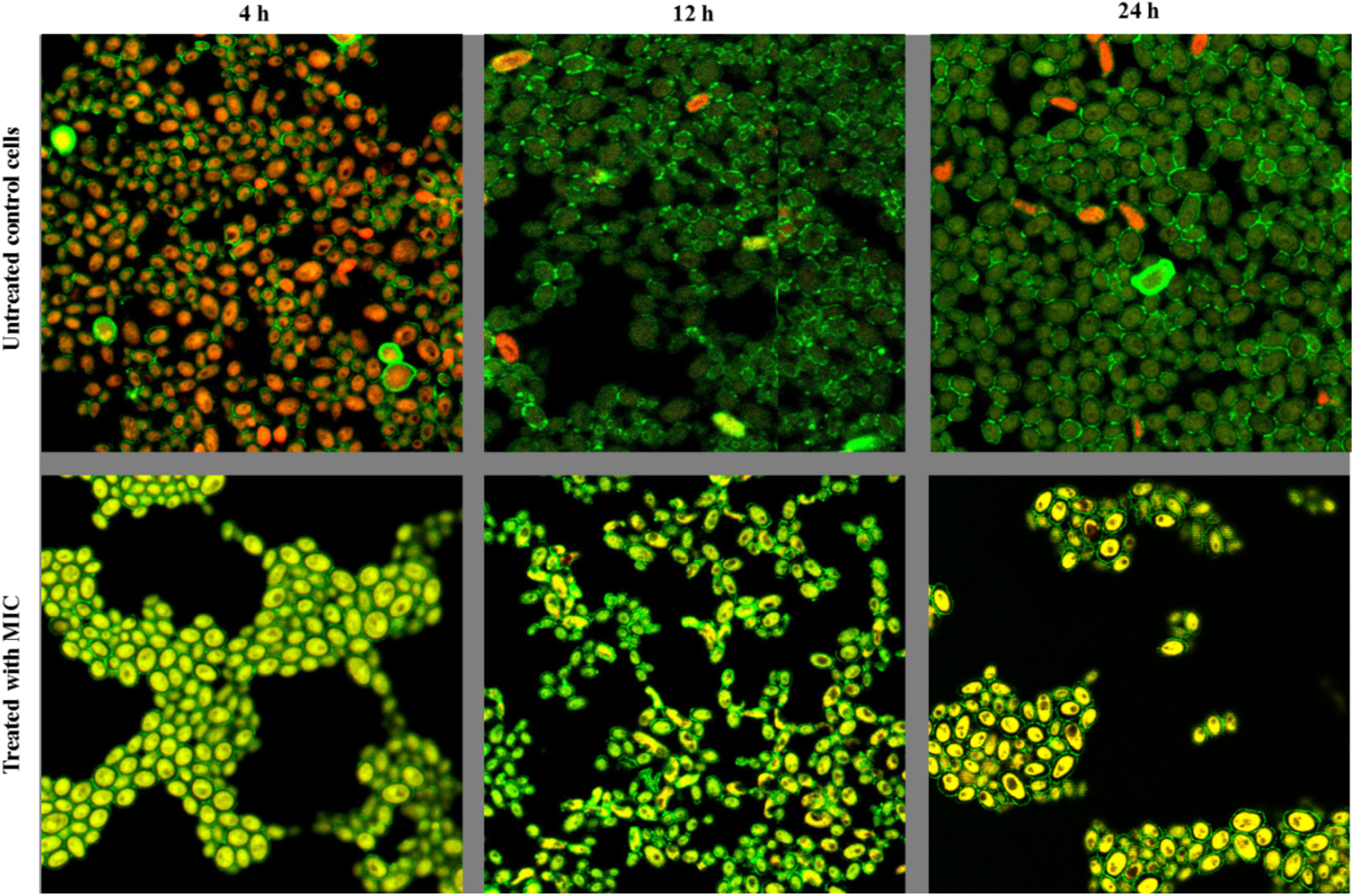
Inhibitory effect of farnesol on *C. auris* biofilms. CLSM analysis of *C. auris* strain MRL 5765 biofilms treated with farnesol at different stages of growth and development (4 h, 12 h and 24 h). The image utilizes ConA-Alexa Fluor 488 conjugate and FUN-1 staining, a 63X oil immersion objective with magnification, X2.5. In above images structures shown in green (ConA) represent the fungal cell wall and biofilm matrix; structures in yellow-green (FUN-1) are metabolically inactive and dead cells; structures in orange-red (FUN-1) are the viable cells.

### Farnesol inhibits R6G efflux mediated transporters

MDR efflux pumps contributes to drug resistance and their functional activity *in vitro* can be studied by using R6G, which is a recognized efflux substrate (34). In fluconazole non-responsive *Candida* cells ATP-dependent efflux pumps (ABC superfamily) throws out R6G dye after entering cells passively (35). In this study, we are measuring the inhibitory/modulatory outcome on MDR efflux transporters of *C. auris*. Here we investigated four isolates of *C. auris* (MRL 2397, MRL 5765, MRL 5762, and MRL 6057) for the passage of substrate, R6G (10µM) after manifestation of farnesol (0.5 × MIC and MIC). When the cells were incubated with PBS (without glucose) we noted an instant uptake of R6G dye by the cells, which came to stability after 30 min. After addition of glucose, control *Candida* cells (untreated) showed energy and time dependent efflux of R6G dye from the cells as they are rich in membrane bound MDR transporter pumps. This was clearly evident from steady rise in the concentration of R6G dye extracellularly (Fig 4). Whereas, farnesol (0.5 × MIC) resulted in inhibition of energy dependent R6G efflux from treated cells confirming its role in modulating/inhibiting MDR efflux transporters. Higher concentration of farnesol (MIC) also had complete inhibitory effect on these energy dependent efflux pumps.

**Fig. 4:**
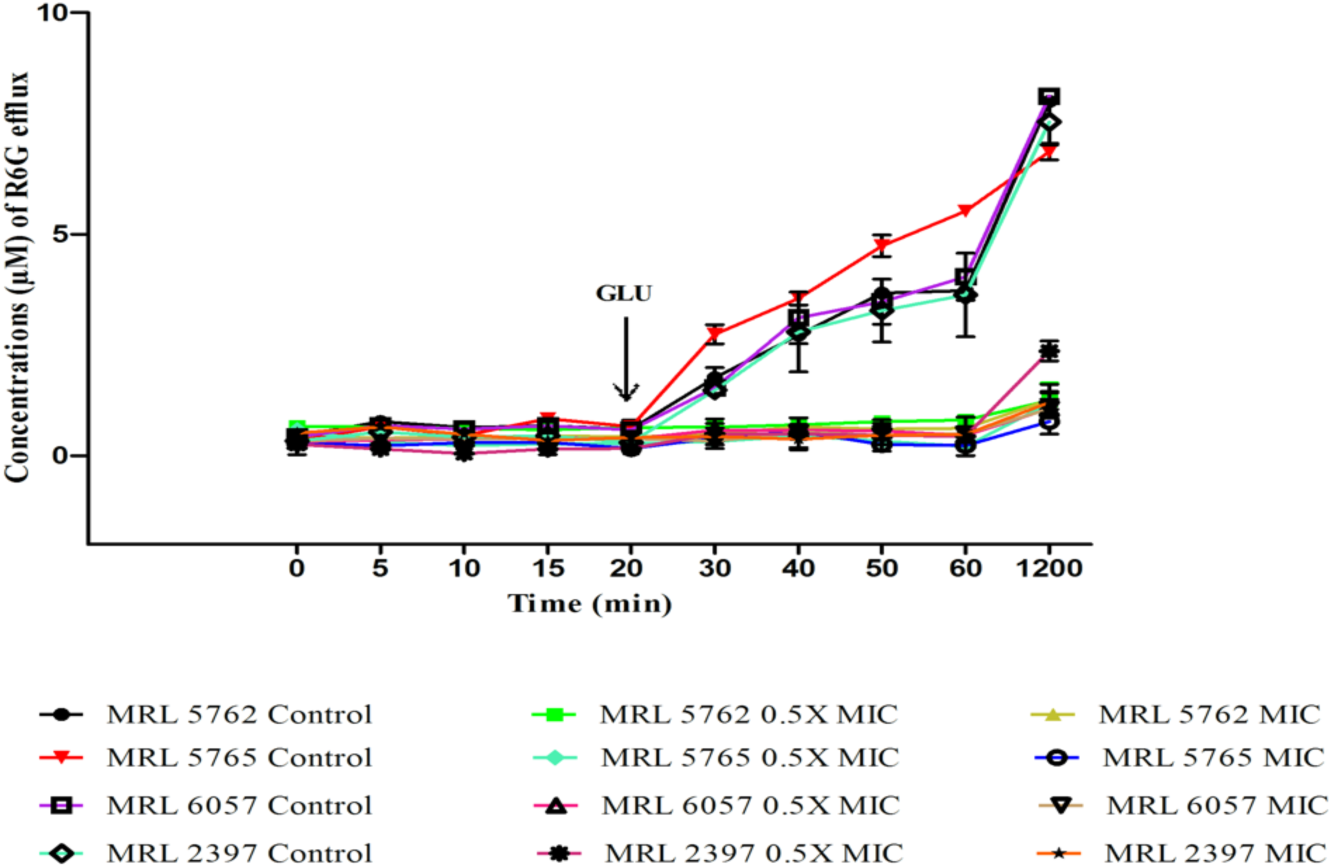
Effect of farnesol on extracellular R6G efflux. Here we are showing concentration of extracellular R6G in untreated control and treated *C. auris* cells. Addition of glucose (0.1M; indicated by an arrow) activated R6G transport from the cells and absorbance was recorded at 527 nm. Concentration of extracellular R6G was determined by standard curve. The values presented here is average of three independent experiments.

Inhibition of efflux pumps in *C. albicans* by phytochemicals has been well established by the researchers (18, 36). Researchers have also explained the role of MDR efflux transporters (ABC and MFS transporter) in resistance and the genes encoding these pumps in *C. auris* (32). Our results are in agreement with previous studies (37), who reported high activity of ABC-type efflux in *C. auris* than *C. glabrata*, isolates proposing *C. auris* with efflux-mediated intrinsic resistance to azoles. It has been reported by authors that in *C. albicans* farnesol particularly targets CaCdr1p and CaCdr2p (ABC transporters), resulting in changes in resistance to azoles in *C. albicans* isolates but in a concentration dependent manner (28). Interestingly, kinetics data cleared that farnesol results in competitive inhibition with an unchanged V_max_ and higher K_m_ values in cells overexpressing CaCdr1p (13).

To the best of our knowledge we are exploring outcome of farnesol on MDR efflux transporters in *C. auris* isolates for the first time. Our results clearly state that farnesol is completely inhibiting ABC MDR transporters similar to that reported in *C. albicans*.

### Farnesol inhibits R6G extrusion

To further confirm the effect of farnesol on activity of *C. auris* efflux pumps, accumulation of R6G dye inside the cells was studied. Untreated *C. auris* isolates were used as a control for estimating actual accumulation of R6G in cells. The fluorescence was recorded high in case of farnesol (0.5 × MIC and MIC) treated *C. auris* cells, suggesting more R6G accumulation intracellularly in comparison with untreated control cells (Fig. 5).

**Fig. 5:**
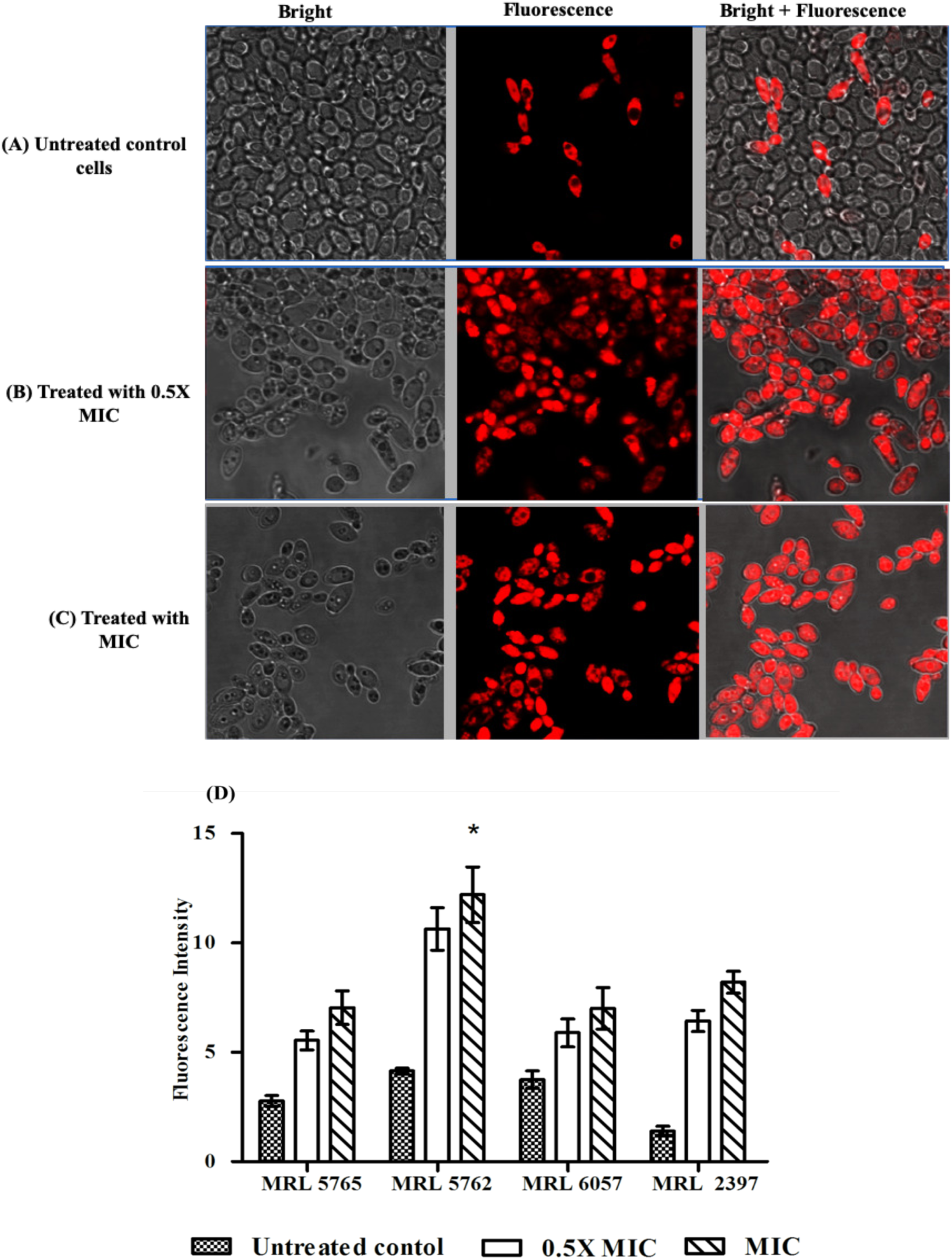
R6G intracellular accumulation. Fluorescence images of R6G stained *C. auris* MRL 5765 cells. The isolate was observed under bright field and fluorescence microscopy. (A) The figure shows untreated *C. auris* cells that have effluxes R6G dye during incubation with glucose and only few cells retain R6G fluorescence. Treatment with 0.5X MIC (B) and MIC (C) results in accumulation of R6G inside the cells after incubation with glucose which is clearly represented by high fluorescence inside the cells. (D) Fluorescence intensity is represented in the form of bar graph and was quantified with the help of Image J software.

*Candida auris* is known to be highly multidrug resistant and among the various resistance mechanisms, the most dominant approach for throwing drug out of cell is the overexpression of MDR transporters and makes *Candida* unresponsive to antifungal therapies. The most common reason for MDR is efflux transporters from ABC superfamily, such as CaCdr1p and CaCdr2p (38, 39). Despite several studies reporting the inhibition of efflux pumps in *C. albicans*, there are no reports on inhibition of efflux transporters in *C. auris* till date. Whereas, lot of work have been published on inhibition of this transporters in *C. albicans* and it has been published that farnesol effectively inhibits/modulates ABC MDR pumps (CaCdr1p and CaCdr2p) and did not affect much in case of MFS efflux pumps (CaMdr1p) (13, 28). This study for the first-time reported efficacy of farnesol to block efflux pumps in *C. auris* isolates.

### Farnesol results in reduced expression of genes associated to resistance

*Candida auris* is an emerging MDR pathogen with higher MIC values against commonly used antifungal drugs when compared to other *Candida* species and the molecular basis of its MDR still remains unclear. Few studies have come up recently that has provided insight into transcriptomic assembly of *C. auris* and revealed the possible mechanism of drug resistance among these isolates (32, 19). For understanding the molecular basis of farnesol on *C. auris* drug efflux pumps and biofilm formation, we studied gene expression in *C. auris. IIF4, PGA26, PGA7, PGA52* and *HYR3* genes played an active role in adhesion of biofilms and *SNQ2, CDR1, CDR2, MDR1*, and *MDR2* genes were functionally related to efflux pumps in *C. auris* (32). After the active role of farnesol on the biofilm and efflux pumps, we studied the effect of farnesol on the expression levels of these genes in different *C. auris* isolates. After treatment with 125 mM farnesol, all the above-mentioned genes were significantly downregulated at varying levels (Fig. 6). Based on these results, we therefore speculated that the inhibition of drug efflux pumps in all the tested isolates by farnesol may be related to downregulation of genes related to efflux pump in *C. auris* (*CDR1, CDR2* and *SNQ2*). The downregulation of *CDR1* and *CDR2* genes after treatment with farnesol confirms its potential to inhibit ABC drug efflux transporters. *MDR1* and *MDR2*, on the other hand, were not having consistent expression level in all the isolates of *C. auris*. It has already been mentioned (19) that *CDR1* is relatively more expressed as compared to *MDR1*. Additionally, both *CDR2* and *MDR2* expression were not raised in any of the isolates. However, in our study both *CDR1* and *CDR2* are over expressed than *MDR1* and *MDR2*. In addition, downregulation of genes (*IIF4, PGA26, PGA7* and *PGA52*) associated to adhesion also supports our *in vitro* biofilm inhibition results. Interestingly, *ALS5* did not expressed in any of the four isolates, which could be due to its late upregulation in the latter stages of mature biofilms or due to ALS-independent adherence mechanism in *C. auris* (32). However, further in-depth studies are required to prove these claims.

**Fig. 6:**
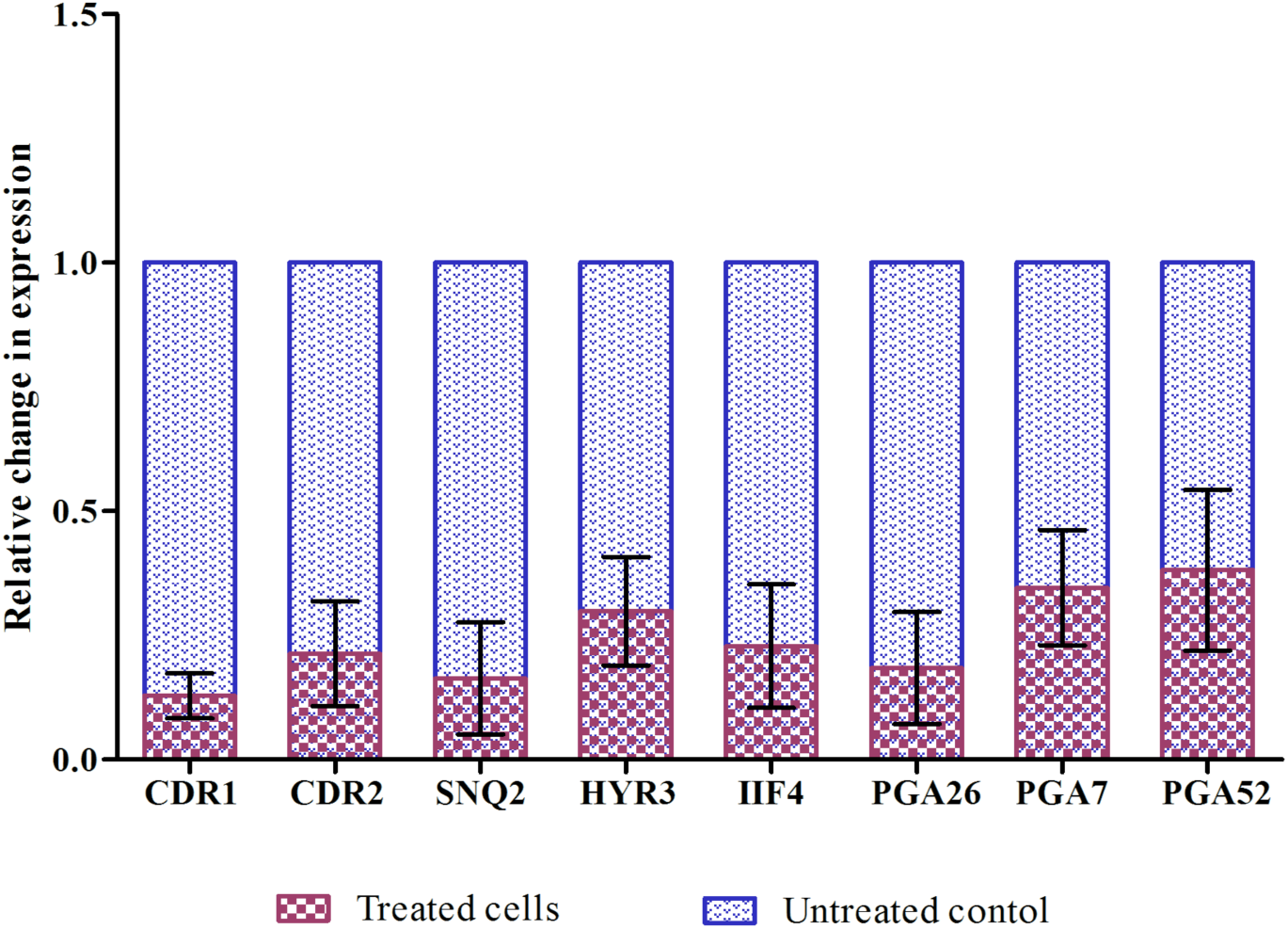
Comparative variation in gene expression of *C. auris* isolates, CDR1, CDR2, SNQ2, HYR3, IIF4, PGA26, PGA7 and PGA52 determined by qPCR. Data represents fold change in gene expression by adopting comparative Ct method. The gene expression between control and treated were compared and was normalized to a value of 1 and ACT1 was used as housekeeping gene.

## Conclusions

Our results showed that farnesol, a quorum sensing molecule, has ability to inhibit growth of multidrug resistant *C. auris* isolates and also has capacity to suppress *C. auris* biofilm formation and efflux pumps. In addition, the effect of farnesol was related to the molecular mechanisms as all the biofilm and efflux pump associated genes were downregulated at varying levels. Furthermore, farnesol has a property to modulate the function of ABC efflux transporters of *C. auris* and thereby can be utilized in reversing azole drug resistance in these isolates. To the best of our knowledge, this is the first study exploring modulatory outcome of farnesol on development of biofilms and efflux pumps of *C. auris* globally and specifically to South African clade, it will be helpful in unrevealing possible mechanisms of action of farnesol against biofilm formation and efflux transporters of *C. auris*

## Author contribution

Conceived and designed the experiments: AA VS. Performed the experiments: VS. Analyzed the data: VS AA. Contributed reagents/materials/analysis tools: AA. Wrote the paper: VS.

## Data Availability

The data that support the findings of this study are available from the corresponding author upon request.

